# Single depolymerizing and transport kinesins stabilize microtubule ends

**DOI:** 10.1101/2020.10.05.326330

**Authors:** Alexandra Ciorîtă, Michael Bugiel, Swathi Sudhakar, Erik Schäffer, Anita Jannasch

## Abstract

Microtubules are highly dynamic cellular filaments and many intracellular processes like cell division depend on an accurate control of their length. Among other factors, microtubule length is actively modulated by motors from the kinesin superfamily. For example, yeast kinesin-8, Kip3, depolymerizes microtubules in a collective manner by a force- and length-dependent mechanism. However, whether single motors depolymerize or stabilize microtubule ends is unclear. Here, using interference reflection microscopy, we measured the influence of single kinesin motors on the stability of microtubules in an *in vitro* assay. Surprisingly, using unlabeled, stabilized microtubules, we found that both single kinesin-8 and non-depolymerizing kinesin-1 transport motors stabilized microtubule ends further by reducing the spontaneous microtubule depolymerization rate. Since we observed this effect for two very different kinesins, it implies a more general stabilization mechanism. For Kip3, this behavior is contrary to the collective force-dependent depolymerization activity of multiple motors. The complex, concentration-dependent interaction with microtubule ends provides new insights into the molecular mechanism of kinesin-8 and its regulatory function of microtubule length.

## STATEMENT OF SIGNIFICANCE

Kinesin-8 motors are important for microtubule length regulation and are over-expressed in colorectal cancer. The budding yeast kinesin-8, Kip3, shortens microtubules collectively. Whether single motors can also depolymerize microtubules remains unclear. Here, we measured the shrinkage speed of microtubules in the presence of different kinesins using interference reflection microscopy. Surprisingly, we found that single Kip3 motors stabilized microtubule ends. Furthermore, even conventional kinesin-1 that transports vesicles had a stabilizing effect implying the presence of a more general stabilization mechanism. A better understanding of kinesin-8 has implications for cell division and associated diseases.

Kinesin motors are involved in many different cellular processes like cell division or intracellular transport (1). They typically translocate along cytoskeletal microtubules and generate force and directed motion, driven by ATP hydrolysis (2, 3). Several kinesins affect the microtubule network itself also during mitosis, which makes them potential targets for cancer treatment (4, 5). For example, members of the kinesin-8 family are known to regulate microtubule dynamics (6–11). The budding yeast kinesin-8 *Kip3* has been shown to depolymerize microtubules in a collective, length- and force-dependent manner (7–9, 12). For depolymerization, it is essential that Kip3 reaches the microtubule end, which requires taking many steps before dissociating from the microtubule. Indeed, Kip3 has a very high run length (7). This high run length originates from a weakly-bound slip state (13) in addition to a second microtubule binding site located in the tail domain (14). This binding site also enhances Kip3’s extraordinary long residence time at the microtubule end and enables Kip3 to crosslink and slide between different microtubules (14, 15). Furthermore, in contrast to its depolymerization activity at high concentrations, it has been suggested that Kip3 stabilizes microtubules at low concentrations (14). This suggestion is motivated by (i) longer microtubules in cells with kinesin-1 mutants that had been fused to Kip3’s tail domain and (ii) *in vitro* assays with microtubules that showed lower shrinkage rates with added Kip3 tail domains. However, direct experimental evidence of microtubule stabilization by full-length Kip3 is missing so far. Moreover, without ATP, rigor-bound conventional kinesin-1 stabilized GDP microtubules (16). Whether this effect persists in the presence of ATP and with tubulin in the GTP state at microtubule ends is also unclear.

To test whether motile kinesin motors can stabilize microtubules, we measured the *in vitro* depolymerization speed of GMPCPP-stabilized microtubules (17) in the presence of different kinesins as a function of their concentration at physiological ATP conditions (Fig. 1, Supplementary Materials and Methods). We used the full-length budding yeast kinesin-8, Kip3, and a truncated rat kinesin-1, rK430, from here on referred to as kinesin-8 and kinesin-1, respectively (Supplementary Materials). Since depolymerization speeds were small, we performed microtubule length measurements over a long period (60 min) using interference reflection microscopy (IRM) (18). IRM enabled label-free microtubule imaging with no significant photodamage and little drift in our setup. In IRM images, surface-immobilized microtubules appear dark (Fig. 1A). We measured their depolymerization speed as the total length change—the difference between the microtubule length in the first and last recorded image—divided by the total acquisition time. Representative kymographs are shown in Fig. 1A. For GMPCPP-stabilized microtubules and in the absence of any kinesin, we measured a spontaneous depolymerization speed of 0.236 ± 0.009 nm/s (mean ± SEM, *N* = 151) corresponding to about 0.88 µm/h (Fig. 1B). At very small kinesin-8 concentrations of 0.7 pM, the microtubule depolymerization speed did not significantly differ from the value without kinesins. For intermediate kinesin-8 concentrations of around 7 pM, the depolymerization speed was reduced about two-fold to 0.11 ± 0.01 nm/s. At higher kinesin-8 concentrations, depolymerization speeds increased again to values that are significantly higher than the one in the absence of kinesin-8. At the highest tested kinesin-8 concentration of 7 nM, we observed the expected length-dependent depolymerization (Fig. 1A, right kymograph showing a non-linear length change with time) in quantitative agreement with the literature (9). Thus, the significant reduction of depolymerization speed at an intermediate number of kinesin-8 motors relative to the spontaneous shrinkage speed, implies a stabilization of the microtubule ends.

**Figure 1.**
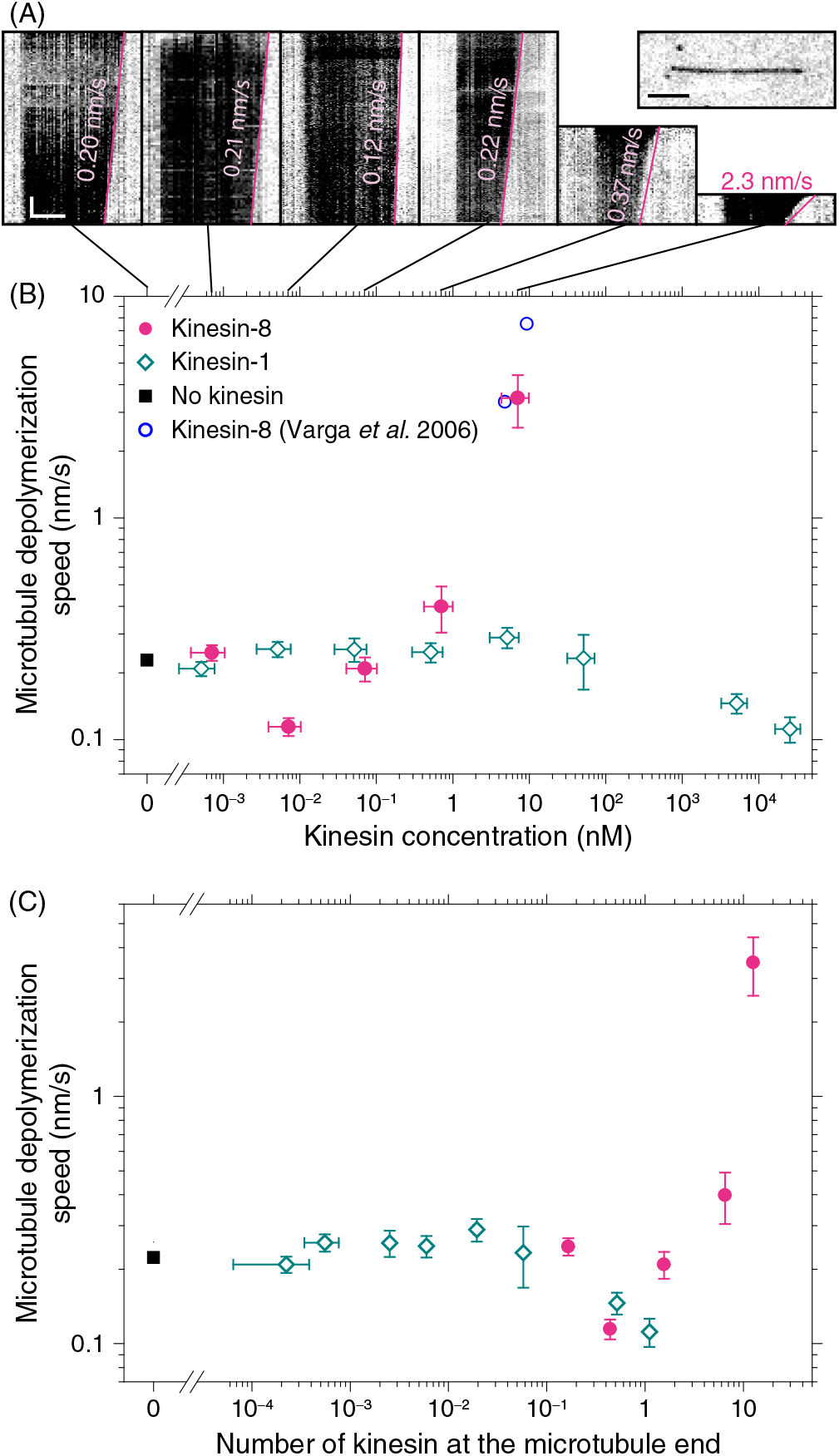
Kinesin stabilizes microtubules. (*A*) Kymographs of microtubules imaged with interference reflection microscopy (IRM) at different kinesin-8 concentrations. The vertical and horizontal scale bar are 10 min and 2 µm, respectively. *Magenta lines* illustrate the shrinkage of microtubules with respective plus-end depolymerization speeds. Upper right-hand corner: Exemplary IRM image of a microtubule (scale bar: 2 µm). (*B*) Total microtubule depolymerization speed as a function of motor concentration for kinesin-8 (*full magenta circles*), kinesin-1 (*open green diamonds*), and in the absence of kinesins (*full black square*). The *open blue circle* corresponds to literature data (7). (*C*) Depolymerization speed data as a function of average number of motors at the microtubule end (see Supporting Material for details).

Do other kinesins stabilize microtubule ends as well? To address this question, we measured the microtubule depolymerization speed at different concentrations of kinesin-1, which does not dwell at microtubule ends, but quickly falls off once the end had been reached. We observed no effect on the microtubule stability for kinesin-1 concentrations below 100 nM (ANOVA, *F* (5, 160) = 0.98, *p* = 0.43). Importantly, for these concentrations, the overall weighted mean of 0.24 ± 0.01 nm/s was not different from the spontaneous microtubule depolymerization speed (*t*-test, *p* = 0.7). However, for very high motor concentrations of 5 µM and 25 µM, we observed a significant decrease of the microtubule depolymerization speed (*t*-test between weighted mean for data below 100 nM and the data point at 25 µM, *p* = 1.8 · 10^−4^). Thus, also a transport kinesin can stabilize microtubule ends at sufficiently high concentrations raising the question of how many kinesins need to be simultaneously at the microtubule end for stabilization.

To determine the average number of kinesins at the microtubule end, we programmed a simulation based on the known motility parameters of the motors (see Supporting Material). Directly measuring the number of kinesins at the microtubule end was not reliable because of insufficient localization precision at the end, photo bleaching during long measurements, or too many fluorophores at higher motor concentrations. Instead, we measured the landing rate as a function of motor concentration (Fig. S1) and used the known translocation speed, run length, and end residence time of the motors to determine for each concentration the average number of motors at the microtubule end. With the relation between the motor concentration and number at the end, we rescaled the concentration axis for both motors (Fig. 1C). Interestingly, for both motors, the stabilization effect occurred roughly at the same number of about one motor per microtubule end. Depolymerization by kinesin-8 occurred at higher numbers that result in traffic jams on individual protofilaments (Fig. S2) consistent with a collective depolymerization mechanism.

In summary, we have shown that if there is on average about one motile kinesin always present at a microtubule end, microtubules are stabilized. Due to kinesin-1’s much smaller run length and end residence time, the concentration at which this condition is met is much higher in comparison to kinesin-8. Surprisingly, already one motor at the end seems to be sufficient to stabilize all protofilaments narrowing down the previous estimate of less than six motors for strongly bound kinesin-1 on GDP-microtubules (16). Our kinesin-8 data directly confirms previous indirect measurements (14) that at low motor concentrations—corresponding to a single motor at the end according to our data—microtubules are stabilized. This finding widens the model of its biological role of microtubule length control. Cells may switch between microtubule stabilization and depolymerization modes by regulating the level of kinesin-8 expression during the cell cycle. Furthermore, since the number of motors at the end scales with microtubule length ((7, 9) and Supporting Material), short microtubules could be stabilized whereas long microtubules may be depolymerized at the same time.

What is the origin of microtubule stabilization by kinesin motors? Previous research suggested inter-protofilament crosslinking via the additional microtubule binding site in the tail domain of kinesin-8 (14). For kinesin-1, a similar electrostatic stabilization could be present through a weak interaction of the neck coiled-coil with the E-hooks of tubulin (19). Another contribution may be from the strongly bound state of kinesin motors at the microtubule end (16, 20), in case of kinesin-1 causing a conformational change of tubulin with an associated elongation and straightening of GDP-tubulin (16). We propose that in addition to these effects, the dimeric state of kinesin links the terminal two tubulin dimers together conferring stability. Recent experiments suggest that both kinesin heads always remain in direct contact with the microtubule lattice during stepping (2, 3). Thus, if a terminal tubulin dimer bound to a kinesin should dissociate from the microtubule lattice, it will still be tethered to the microtubule end via the kinesin dimer. In this manner, this terminal tubulin may quickly reattach with an effectively increased on-rate. This increase would lead to an overall stabilization of the microtubule end. Because of the three-start-helix structure of the microtubule (1) and assuming that the microtubule end is blunt (17), there is only one tubulin dimer at the very end of the helix that has only one lateral tubulin contact. Therefore, already one kinesin might be sufficient to stabilize this “weakest” spot of the microtubule. Further experiments with other kinesin dimers and possibly with truncated tail domains or modified neck coiled coils may be necessary to establish a general stabilization mechanism of kinesin dimers and a possible contribution from further microtubule interactions of their tails. While for kinesin-8 microtubule stabilization at low concentrations may be biologically relevant, we do not expect a relevant *in vivo* effect for kinesin-1 since unnaturally high motor concentrations were required. For kinesin-8, our observations support and widen the model on the dual-mode regulation of microtubule dynamics by this kinesin.

## SUPPORTING MATERIAL

Supporting Methods and Material and two figures are available at http://www.biophysj.org/biophysj/supplemental/XXX. Single kinesins stabilize microtubules

## AUTHOR CONTRIBUTIONS

A.J. and E.S. designed research; A.C., A.J., and S.S. performed measurements and analyzed the data. M.B. wrote the simulation and analyzed its results. M.B., A.J., and E.S. wrote the manuscript. All authors commented on the manuscript.

## ACKNOWLEDGMENTS

This work was supported by the German Research Foundation (DFG, JA 2589/1-1), the interdisciplinary “nanoBCP-Lab” funded by the Carl Zeiss Foundation (Forschungsstrukturprogramm 2017), and the PhD Network “Novel Nanoparticles” of the Universität Tübingen. Alexandra Ciorît,ă was co-funded by the Erasmus+ programme of the European Union and by the Romanian Ministry of Research and Innovation (CCCDI – UEFISCDI, project number PN-III-P1-1.2-PCCDI-2017-0387 / 80PCCDI / 2018, within PNCDI III). We would like to thank Andreas Neusch and Katrin Luginsland for preliminary experiments. We thank Viktoria Wedler and Carolina Carrasco-Pulido for comments on the manuscript.

## SUPPORTING CITATIONS

References (21–39) appear in the Supporting Material.

