## Supporting Material for "Single depolymerizing and transport kinesins stabilize microtubule ends"

### Single transport and depolymerizing kinesin stabilize microtubules

#### S1 MATERIALS AND METHODS

##### S1.1 Sample preparation and assay

Porcine tubulin (3 µM) was polymerized in PEM buffer (80 mM PIPES, 1 mM EGTA, 1 mM $MgCl_2$, pH = 6.9) with 1 mM $MgCl_2$ and 1 mM guanosine-5'-[($\alpha$,$\beta$)-methyleno] triphosphate (GMPCPP, Jena Bioscience, Jena, Germany) for 1.5 h at 37 °C. Afterwards, the GMPCPP-stabilized microtubules were diluted with PEM, spun down in a Beckman airfuge, and resuspended in PEM.

Full-length budding yeast kinesin-8 (Kip3-eGFP-$His_6$) was expressed and purified according to (1, 2). The motility buffer for kinesin-8 stepping assays was PEM supplemented with 1 mM ATP, 0.1 mg/ml casein, 112.5 mM KCl, and a reducing agent and oxygen-scavenger system (10 mM DTT, 20 mM glucose, 20 µg/ml glucose oxidase, 8 µg/ml catalase) (1, 3–5). Truncated rat kinesin-1 (rK430-eGFP-$His_6$) was expressed and purified according to (6). The stock concentration of 51 µM limited the maximum concentrations we could test in the assays. The motility buffer for kinesin-1 stepping assays was the same as for kinesin-8 except no KCl was added.

Experiments were performed in flow cells that were either constructed using silanized, hydrophobic glass cover slides or supported solid lipid bilayers slides as described before (3, 7). The depolymerization speed and the kinesin landing rate did not significantly differ for both glass treatment. The channels of flow cells were washed with PEM, filled and incubated successively with anti $\beta$-tubulin I (monoclonal antibody SAP.4G5 from Sigma in PEM), Pluronic F-127 (1 % in PEM), and GMPCPP-stabilized microtubules in PEM (1, 3–5). After 10 min of microtubule incubation, different kinesin dilutions in motility buffer were flown in and the measurements were started. For control measurements without kinesin, only motility buffer was used.

##### S1.2 Data acquisition and photo damage

To enable long term measurements, microtubules were visualized using a custom-built interference-reflection-microscopy (IRM) setup with a light-emitting diode (LED, center wavelength: 450 nm) as a light source (8, 9). The microscope is combined with simultaneous total internal reflection fluorescence (TIRF) microscopy (2, 8) and equipped with a millikelvin-precision temperature control (10) set to 30.000 °C. IRM images were acquired at 25 fps and averaged over 100 frames resulting in a final acquisition speed of 0.25 fps. The lateral and axial drift was at most 0.3 nm/s and 0.5 nm/s, respectively. The axial drift was manually corrected during the measurements.

To test whether the LED light had an influence on the microtubule depolymerization speed, we measured the microtubule length change in control experiments by taking only one image in the beginning, switching the LED off, and then, after 60 min, switching the LED on again to take one final image. As expected based on previous measurements (5), the LED light had no statistically significant effect on the microtubule depolymerization speed. Thus, the LED was always switched on.

To measure the landing rate of kinesin on microtubules, TIRF measurements were performed after each IRM measurement (data not shown). Kinesin motility was visualized with time-lapse videos recorded at 1 fps for kinesin-8 and 10 fps for kinesin-1 with an exposure time of 100 ms. The laser output was set to 10 mW (488 nm).

#### S1.3 Data and statistical analysis

The IRM and TIRF images were processed and analyzed using the software Fiji (11). IRM image stacks were contrast and brightness adjusted, and corrected for lateral drift (12, 13). Kymographs were generated by a custom written macro.

To analyze the microtubule depolymerization speed, we measured the total length change of single microtubules over 60 min by taking the first and last IRM image. Due to the collective depolymerization activity of kinesin-8, at higher concentration (0.7 nM, 7.1 nM) the acquisition time was reduced to less than 20 min. The average microtubule length was 8.3 ± 3.9 µm (mean±SD, $N$ = 610). We only analyzed microtubules that were longer than 3 µm. For the different kinesin-1 and kinesin-8 concentrations, we analyzed in total 195 and 264 microtubules, respectively. As a control, the spontaneous depolymerization speed of 151 microtubules without kinesins was analyzed. We did not observe a significant difference in depolymerization speeds for the two motility buffers.

As the microtubule polarity cannot be observed in IRM images, we measured the total microtubule length change. Thus, the calculated depolymerization speed corresponds to total microtubule depolymerization (minus and plus end together). In control measurements, we detected the microtubule polarity by the direction of kinesin motility using TIRF microscopy. The observed spontaneous microtubule minus end depolymerization speed of 0.06 ± 0.03 nm/s (SD, $N$ = 74) was independent of the kinesin-8 concentration (ANOVA, $F(3, 70) = 0.28$, $p = 0.84$).

We estimate the precision for a single microtubule length measurement to be about 2 px corresponding to 210 nm. Thus, the measurement error for the length difference between the last and first image of a single microtubule is $l_{\mathrm{err}} = \sqrt{2} \cdot 210\,\mathrm{nm} \approx 300\,\mathrm{nm}$ resulting in a measurement error for the depolymerization speed of a single microtubule during the measurement time $t_{\mathrm{msr}}$ to $v_{\mathrm{err}} = l_{\mathrm{err}}/t_{\mathrm{msr}} = 300\,\mathrm{nm}/3600\,\mathrm{s}$ = 0.08 nm/s. With roughly 40 microtubules per concentration, the depolymerization-speed measurement precision was about 0.01 nm/s.

In the absence of kinesin-8 and for small kinesin-8 concentrations (0.7 pM, 7 pM, and 0.07 nM), the microtubule depolymerization speed did not correlate with the microtubule length (Pearson correlation coefficients: 0.1, 0.4, 0.1 and 0.3, respectively). For higher kinesin-8 concentrations, we observed the expected microtubule length-dependent depolymerization. As we analyzed the total length change over the measurement time, our depolymerization speed is underestimated but still in good agreement with reported speeds (14).

Statistical tests were performed using two-tailed, unpaired $t$-test and one-way ANOVA with a confidence level of $\alpha$ = 0.05.

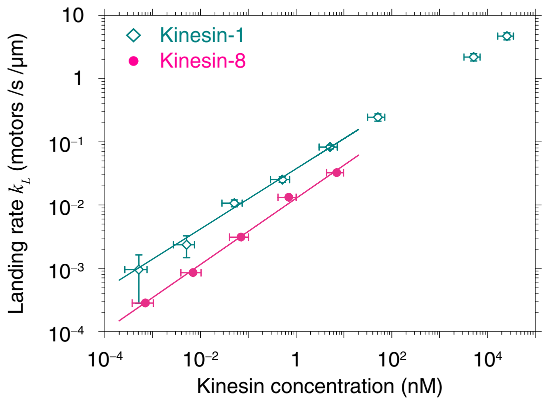

FIGURE S1 Kinesin landing rate as a function of concentration for kinesin-8 (*solid magenta circles*) and kinesin-1 (*open green diamonds*). Error bars correspond to propagated errors due to the Poisson counting error of landing events. The *magenta* and *green lines* refer to power-law fits with exponents of 0.52 ± 0.03 and 0.48 ± 0.02, respectively (standard errors). The landing rate values for the three highest kinesin-1 concentrations were extrapolated using the fit.

### S2 KINESIN LANDING RATE

To determine the number of kinesin motors at the microtubule end as a function of motor concentration, we measured the landing rates as an input for motility simulations (see Sect. S3). We determined the motor landing rate by counting the landing events (a motor localization on microtubules for at least three subsequent frames) and divided this number by the measured total microtubule length and measurement time (Fig. S1). For the three highest kinesin-1 concentrations, for which single motor traces could not be distinguished any more, we extrapolated the landing rate based on a power-law fit to the landing rate measured for small motor concentrations. Interestingly, the measured landing rate for both kinesins scaled with the square root of the kinesin concentration contrary to the commonly assumed linear relationship. One possible explanation for the square-root scaling might be a competitive tetramerization reaction of the kinesin motors with increasing concentration assuming that tetramers are inefficient in landing and motility. Oligomerization might be due to the eGFP tags that have been shown to dimerize (15).

### S3 KINESIN-AT-MICROTUBULE-END SIMULATION

Since the measurement of the number of kinesins at the microtubule end was not reliable, we simulated the kinesin motility on microtubules. Modeling the behavior of motors has been a useful tool to determine kinetic motor parameters or collective multi-motor behavior based on measured motility parameters of single motors (see e.g. (3, 16) or (17–20), respectively). Our Monte-Carlo simulation was implemented in MATLAB (MathWorks). A single microtubule was represented by a Boolean matrix with $N_{PF}$ rows for the different protofilaments and $N_x$ columns for the kinesin binding sites along the protofilaments. We used $N_{PF} = 14$ for GMPCPP-stabilized microtubules (21). Occupied sites were set to *true* and unoccupied ones to *false*. Motor landing, stepping, or detachment could occur during consecutive simulation time steps of duration $dt$, implemented as described by Arpağ *et al.* (22): An unbound motor from the bulk landed on the microtubule with a probability $P_{\text{land}} = k_L L_{\text{MT}} dt$, where $k_L$ is the landing rate and $L_{\text{MT}}$ the microtubule length chosen to roughly match the mean length of our experiments. A landing event happened if a randomly generated number $R$ in the interval $[0, 1]$ fulfilled $R < P_{\text{land}}$. In case of a landing event, the motor occupied a random, free binding site that was set to *true* in the microtubule matrix. Also, this motor was randomly assigned a specific run length out of an exponential distribution with a mean run length $L_R$.

Bound motors could take a forward step of size $\delta$ corresponding to a center-of-mass motor displacement of 8 nm. Sideward or backward steps were not considered. The step dwell time was also exponentially distributed with the step probability $P_{\text{step}} = 1 - \exp(-(\upsilon/\delta)dt)$. If a motor took a step, its current matrix element was set to *false* and the next matrix element (an increment in column number) was set to *true* under the constraint that the next binding site was free. If the next binding site was occupied, no step and also no detachment occurred. If two consecutive sites were occupied, we call this situation a "traffic jam". For all motors, the current run length and time spent on the microtubule lattice were saved. If the current run length exceeded the motors assigned run length, the motor detached from the microtubule lattice resulting in an unoccupied binding site. If a motor reached the microtubule end, a random end residence time out of an exponential distribution with a mean $t_{\text{end}}$ was assigned to the motor. Detachment from the microtubule end occurred when this time was exceeded. Note that a decrease of end residence times by traffic jams or force (5, 23) was not considered here.

The duration of each simulation corresponding to one microtubule is given by the number of iterations times the simulation step time $dt$. The step time $dt$ was sufficiently small compared to the smallest time constant. For each time point, the number of motors in the bulk, on the microtubule lattice, and at the microtubule end were counted with the total number of motors $N_{\text{total}}$ remaining constant. To reduce stochastic noise, we repeated the simulation. In essence, each repetition corresponded to a new microtubule. Since the simulation was started with an empty, unoccupied microtubule, it took some time to reach a steady state. To determine the mean value of motor numbers in a specific location, motor counts as a function of time for a certain number of simulations were averaged. Subsequently, model functions were fitted to the time course as shown in Fig. S2A. Fit results were then used for further data analysis.

To simulate the activity of kinesin-8, we chose the following parameters according to literature values (5, 23–25): speed $\upsilon = 50$ nm/s, run length $L_R = 12$ μm, end residence time

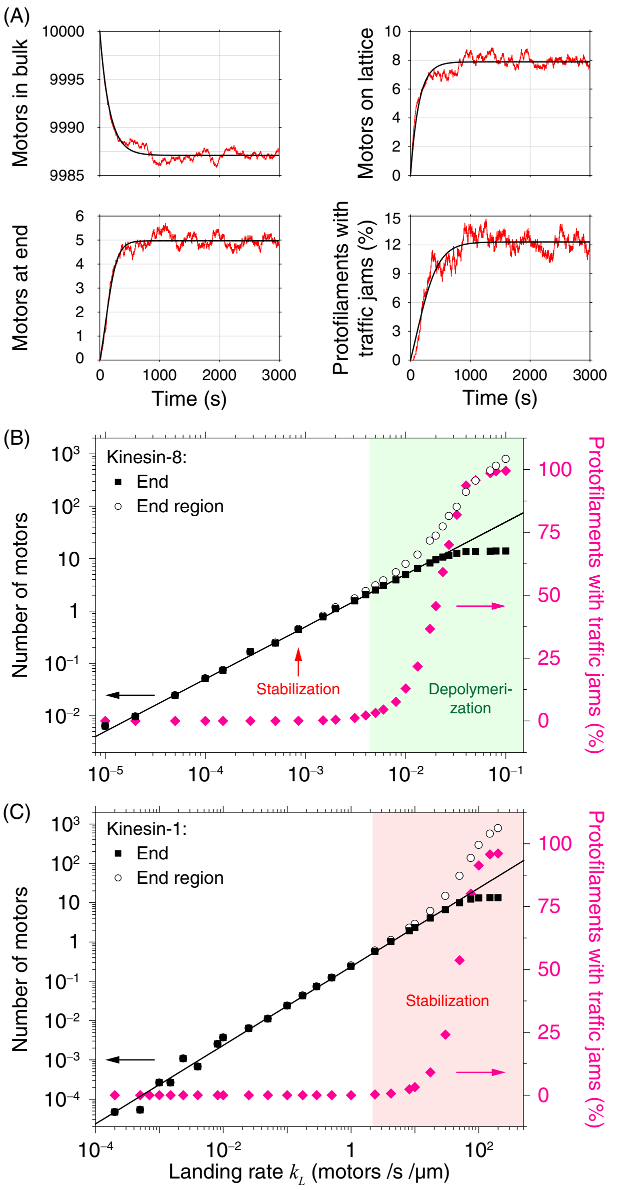

FIGURE S2 Simulation of kinesin motility on microtubules. (*A*) Mean number of motors (*red lines* are averages of 50 simulations) in the bulk, on the microtubule lattice, and at the microtubule end as well as percentage of microtubule protofilaments with traffic jams as a function of time. *Black lines* are fits with exponentials (top) and logistic functions (bottom). (*B,C*) Steady state number of motors at the microtubule end (*solid black squares*), of motors waiting at the end in traffic jams (end region, *open black circles*), and the percentage of protofilaments with traffic jams at the end (*solid magenta diamonds*) as a function of landing rate $k_L$ with parameters for (*B*) kinesin-8, (*C*) kinesin-1. For small landing rates, lines were fitted to the number of motors at the end (*black lines*). The *red arrow* marks the landing rate, at which we measured microtubule stabilization by kinesin-8, i.e. a minimum depolymerization speed. The *green shaded area* marks the concentration range, for which significant depolymerization by kinesin-8 is expected. The *red shaded area* corresponds to concentrations, for which we measured a decreased depolymerization speed by kinesin-1.

$t_{\text{end}} = 86.5$ s, and microtubule length $L_{\text{MT}} = 8$ µm—defining the number of binding site per protofilament $N_x = L_{\text{MT}}/\delta$, and a step time $dt = 10$ ms. To ensure an access of motors in the bulk, we chose a total motor number of $N_{\text{total}} = 10,000$. To simulate different motor concentrations, we varied the landing rate $k_L$ over the range we measured (Fig. S1). In Fig. S2B and C, the simulation results are plotted as a function of landing rate. As expected for small landing rates, the mean number of motors at the microtubule end increased linearly with the landing rate. We used this plot together with Fig. S1, to map the number of motors at the microtubule end $N_{\text{end}}$ to the landing rate $k_L$ and corresponding pipetted motor concentration. With this mapping, we converted the concentration axis of Fig. 1B in the main text to the motor number at the microtubule end plotted in Fig. 1C.

For a kinesin-8 concentration and corresponding landing rate, for which we measured microtubule stabilization, the simulation showed that only about one kinesin-8 was on average at the end (Fig. S2B). At higher landing rates, the motor number at the end saturated and converged to the total number of protofilaments $N_{PF}$, i.e. all protofilament ends were occupied. At kinesin-8 concentrations and corresponding landing rates, for which we measured active depolymerization, the number of motors at the end, the number of motors waiting in traffic jams, and the percentage of protofilaments with traffic jams at the end were significantly higher.

We repeated the simulations as a function of landing rate for kinesin-1 using the following parameters (23, 26, 27): speed $v = 800$ nm/s, run length $L_R = 0.8$ µm, end residence time $t_{\text{end}} = 0.3$ s, and simulation step time $dt = 0.1$–$1$ ms. The results are plotted in Fig. S2C and show that the mean number of motors at the microtubule end saturated only at very high landing rates. At kinesin-1 landing rates, for which we measured microtubule stabilization, the simulation also resulted in about one kinesin-1 at the end, i.e. about the same number as for kinesin-8.

In additional simulations, we found that $N_{\text{end}}$ increased linearly with $t_{\text{end}}$, $L_{\text{MT}}$, and $L_R$ (for $L_R \ll L_{\text{MT}}$), and saturated for $L_R \gg L_{\text{MT}}$ (data not shown). $N_{\text{end}}$ did not depend on the forward speed $v$. Simulations also confirmed that the average number of motors on the microtubule at steady state scaled linearly with the landing rate.

The simulation enabled us to not only count the number of motors at the end but also the number of motors that were waiting at the end in traffic jams and the corresponding percentage of protofilaments with traffic jams at the end (Fig. S2). Both values are important for and consistent with the "bump-off model" of kinesin-8's cooperative microtubule depolymerization. In this model, high numbers of kinesin-8 at the end accelerate microtubule shrinkage (23). In our simulation, we did not consider a decreased residence time of end-bound motors in traffic jams as observed by Varga *et al.* (23). If we would consider this effect, saturation effects would shift to higher landing rates. However, since we were not interested in this effect here, we did not implement it. Overall, the simulation provided quantitative insight into how many motors were located where on the microtubule—information that is difficult to access experimentally and is limited by the resolution and contrast of the experiment.
